# Subconcussive preconditioning prevents microglial morphology changes and improves cognitive outcomes in mice

**DOI:** 10.1101/2025.04.28.651064

**Authors:** Erin D. Anderson, Kyulee Kim, Anastasia P. Georges, Annika Naveen, Elena Grajales, Daunel V. Augustin, David F. Meaney

## Abstract

Subconcussive impacts are highly prevalent in contact sports and are thought to increase concussion risk. However, the specific conditions under which these subconcussive impacts influence concussion outcomes are uncertain, limiting our understanding of the mechanisms behind repetitive head trauma. Given that subconcussive impacts elicit a microglial response, and microglial morphology offers insight into function, we examined how subconcussive preconditioning affects microglial morphology and cognitive outcome after concussion. To investigate this question, we developed and validated a scalable, closed-head controlled cortical impact model. Using this approach, we found that although concussion elicited features of hypersurveillant microglia at 1 day post-injury, they resolve by 9 days post-injury, and subconcussive impacts only produced microglial changes at 9 days post-injury. When subconcussive impacts preceded a concussive impact (i.e., preconditioned concussion) no changes in microglial morphology appeared at either 1 or 9 days after injury. Interestingly, subconcussive preconditioning eliminated concussion-associated cognitive deficits in novel object recognition and this cognitive protection was time dependent: preconditioning impacts were only protective if delivered within 2 minutes of concussion, and had no effect if delivered over a 48-hour window. These results suggest that some types of subconcussive impacts may offer protection against subsequent concussion and mitigate changes in microglial morphology. Understanding this timing window could inform strategies for minimizing cognitive impairments in athletes exposed to repetitive head trauma.

## Introduction

An estimated 1.6-3.8 million concussions occur due to sports- and recreation-related activities each year.^1^ Subconcussive head impacts, defined as impacts that do not produce immediate neurocognitive deficits, can affect the likelihood of sports-related concussion.^2–7^ Although there is a consensus that the hundreds to thousands of subconcussive head impacts that a contact sports player may experience over a season are detrimental to long-term brain health,^8,9^ it remains unclear how these repeated exposures affect concussion outcome.^5–7,10,11^ A persistent challenge in these human studies is uncontrolled variability amongst subjects, including unmeasurable history of head impacts, genetic predisposition, and psychological factors, which collectively impede our precise characterization of how subconcussive impacts influence concussion likelihood.^12–15^ A second challenge is that these human studies preclude a mechanistic understanding of the brain’s cellular response to these repeated exposures, limiting insights that could lead to a more effective treatment of concussion.

Rodent studies, however, afford greater experimental control and can be used to identify important impact features and the cellular processes they modify. In particular, rodent studies show that the number of impacts and the timing of those impacts are key modifiers of outcome within the first month after injury (Fig 1 A). Although it is possible for mice to sustain a few *subconcussive* impacts and remain cognitively and histologically normal, increasing the number of subconcussive impacts, especially over a long period of time, increases the likelihood of concussion-like outcomes, such as in particular when it comes to microgliosis and white matter degeneration.^16–25^ Past studies show that repeated concussions are consistently damaging, but increasing the impact window (the time between first and final impact) lessens their cumulative severity, especially in regard to microgliosis, astrogliosis, and axonal damage (reviewed in Bolton-Hall et al., 2019^26^). Taken together, these studies suggest that a large number of subconcussive impacts, delivered over a long period of time, will likely worsen concussion outcomes, while delivering fewer subconcussive impacts over a long period of time may reduce their cumulative effect. Indeed, Harper and colleagues found that a small number of subconcussive blast injuries delivered over 1-2 days were protective against a concussive blast injury and implicated the kynurenine pathway, which exists at the interaction of neural and immune systems, as a mediating factor.^27^

**Fig 1.**
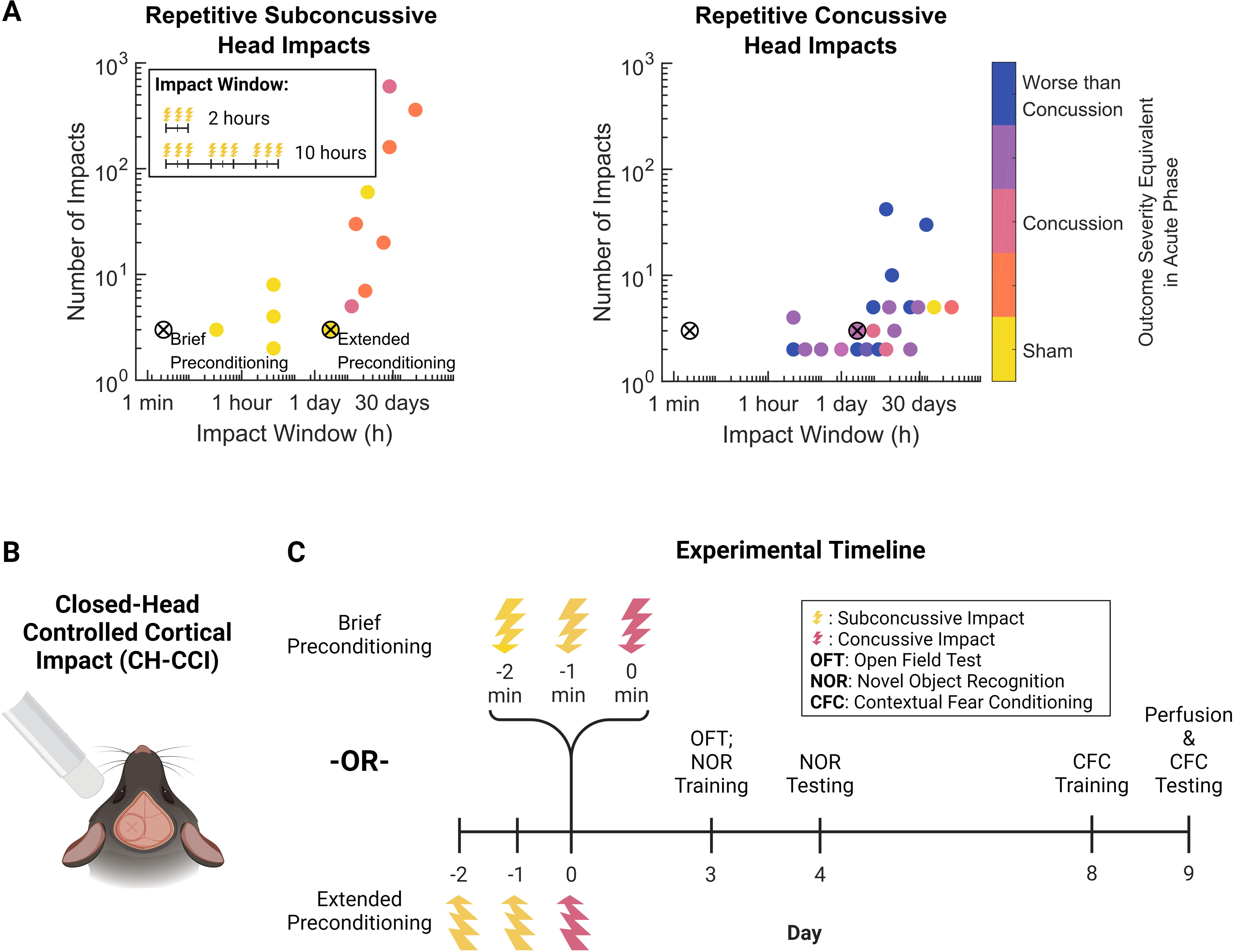
Closed-head controlled cortical impact for subconcussive preconditioning. (A) Review of acute (<1 month) outcomes of repetitive subconcussive ^16–24^ and concussive ^26^ head impacts models in rodents. Impact window is defined as the difference in time between initial and final impact. In our study, brief preconditioning and extended preconditioning are defined as occurring over 2 minutes and 48 hours, respectively. Outcomes are broadly defined to include glial activation, behavioral deficits, and neuronal changes. (B) Rubber tip used to impact the exposed left parietal bone of the animal’s skull. (C) Experimental timeline for impacts, behavioral assessment, and sample collection. Created using BioRender.

Microglia are often studied as potential mediators of repeated concussive and subconcussive impacts, perhaps because of the ability to rapidly activate several innate immune responses in the brain. Repetitive subconcussive impacts induce a microglial response,^19,24^ and even single subconcussive impacts can elicit a microglial response in the absence of behavioral deficits.^28^ Microglia are beneficial to injury recovery by migrating to the site of injury, clearing debris, and coordinating a multicellular injury response through the release of inflammatory molecules (e.g., cytokines). However, persistent microglial activation can be detrimental to injury recovery, leading to neurotoxicity, chronic neuroinflammation, and aberrant synaptic remodeling.^29^ Repetitive microglial stimulation is similarly multifaceted: depending on the nature of the stimulus (either “priming” or “desensitizing”), microglia can exhibit a stronger or weaker reaction as part of innate immune memory.^30,31^ While the precise conditions which govern whether a stimulus is priming (entrains an exaggerated pro-inflammatory immune response) or desensitizing (induces a tolerance which suppresses the immune response) remain unclear, studies using lipopolysaccharide (LPS) as a stimulus suggest that dose strength, duration, timing, and sequence are critical factors influencing outcome.^30,32^ Indeed, preconditioning with LPS can be neuroprotective not only for TBI,^33–35^ but also ischemic brain injury^36^ and epilepsy.^37–40^ Preconditioning using other methods, such as transient ischemia or subthreshold cortical stimulation, can also be neuroprotective against ischemic brain injury and epilepsy and may involve microglia-dependent mechanisms.^41–44^

In the present study, we hypothesized that subconcussive impacts delivered prior to a concussion would modify the microglial response and, in parallel, prevent the cognitive deficits that normally occur after a concussive impact. Using a novel closed-head controlled cortical impact model, we found that microglial morphology was sensitive to impact loading conditions. Brief subconcussive preconditioning desensitized the microglial response, preventing the hyperramified morphology associated with concussion. This desensitization correlated with the prevention of concussion-associated cognitive deficits, but only if preconditioning occurred within minutes of the concussive impact. Extended preconditioning over days eliminated this protective cognitive effect. Our findings show that microglia are one key component in mediating the brain’s response to repetitive head impacts and suggest that carefully timed subconcussive exposures can modulate microglial reactivity, potentially influencing cognitive outcomes after concussion. This work contributes to our understanding of the complex interplay between subconcussive impacts, microglial activation, and cognitive function, with implications for concussion prevention and treatment strategies in sports and other high-risk activities.

## Materials and Methods

### Animals

In this study, we used adult male C57BL\6 mice (8-12 weeks old; Charles River Laboratories). Mice were housed in an animal care facility with a 12h light/dark cycle and *ad libitum* access to food and water. Upon arrival, animals were fed Lab Diet 5001(Animal Specialties and Provisions). All animal procedures were approved by the University of Pennsylvania’s Institutional Animal Care & Use Committee (IACUC).

### Experimental Design

#### Closed-Head Controlled Cortical Impact

For our study, to answer the question of how subconcussive impacts affect concussion outcomes, we wanted to select or develop an injury model that would address some of the challenges in interpretability experienced by human studies. Heterogeneity in the impact number, location, and timing, as well as low concussion incidence, have made it difficult to precisely define how subconcussive impacts affect concussion risk in humans.^12^ To mimic the head impacts experienced by humans, we wanted to use a closed-head impact model. However, no existing model permits consistent control of impact site and severity, two features which would ensure precise preconditioning impacts and subsequent injury over the exact same site. Traditional controlled cortical impact, although accurate and reproducible, requires a craniectomy to deliver impacts.^45^ Weight drop, although closed-head and scalable, has poor accuracy and repeatability.^46^ Closed head impact model of engineered rotational acceleration (CHIMERA) although closed-head and scalable, produces diffuse pathology throughout the entire brain and precludes ipsilateral/contralateral comparisons.^47^

To overcome the shortcomings of existing models for our study’s design, we developed a closed-head impact that we could scale to deliver concussive and subconcussive impacts to one hemisphere. We outfitted the Impact One Controlled Cortical Impact Device (Leica Biosystems) with a custom 6 mm diameter rubber tip (Model # E10BP-K6, Pentel). Prior work suggests that impact severity scales with velocity, and helmeted impact velocities scale between humans and rodents.^48^ For concussive impacts, we injured the mice using an impact velocity of 3.5 m/s, which we confirmed in pilot studies to produce behavioral deficits. We then scaled the impact speed to produce a subconcussive impact, impacting the mice at 1.0 m/s, consistent with reports of subconcussive impacts occurring in humans at 25-33% of the magnitude of concussive impacts.^49,50^ We further confirmed that a single subconcussive impact produced no behavioral deficit, consistent with its clinical definition (Fig 2).

**Fig 2.**
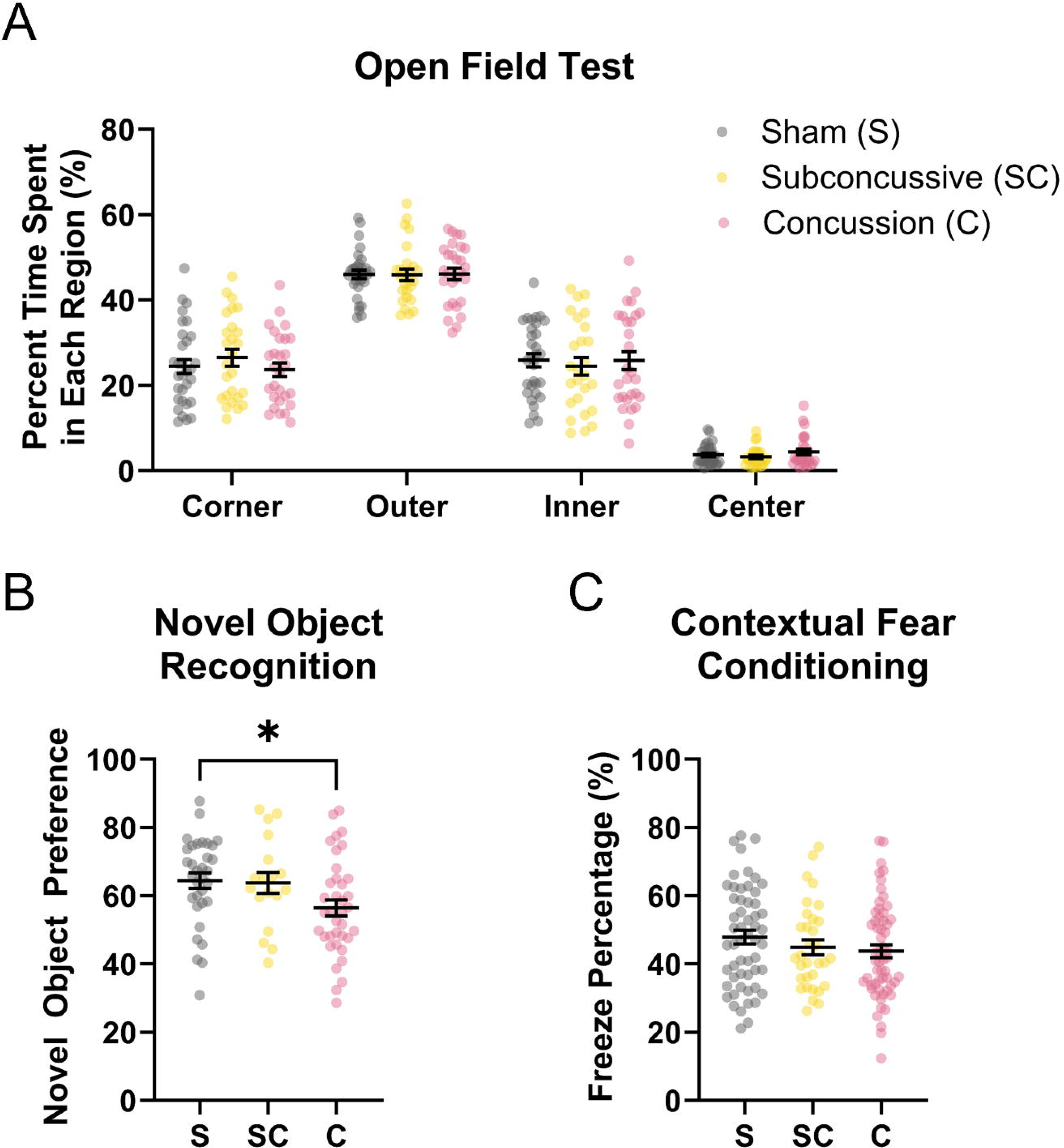
Cognitive impairment scales with injury severity. (A) Neither subconcussive nor concussive impacts have a significant deficit in percent time spent in the corner, outer, inner or center areas of the open field. (B) Concussion exhibits a significant deficit in novel object recognition. (C) Neither subconcussive nor concussive impacts have a significant effect on freeze percentage. * indicates p<0.05. Statistical testing was performed using two-way ANOVA (A) or one-way ANOVA (B, C) with Tukey’s post-hoc test. Data are shown as mean ± SEM. S: Sham; SC: Subconcussive; C: Concussion.

To perform either subconcussive or concussive impacts, we induced anesthesia with 3.5% isoflurane and maintained it with 1.5-2% isoflurane. We then made an approximately 10 mm incision above the midline of the skull and aligned the impactor over the central aspect of the left parietal bone. We set the dwell time to 3 ms and the impact depth to 1 mm to prepare for impact and delivered no impact (sham), a 1 m/s impact (subconcussion), a 3.5 m/s impact (concussion), or two 1 m/s impacts followed by a 3.5 m/s impact (preconditioned concussion). We sacrificed any mouse that experienced skull fracture with bleeding, consistent with Macheda et al., 2022.^51^ We also examined the skulls at the endpoint and excluded any mice for visibly unhealed skull fracture. Unconditioned mice (sham, single subconcussion, single concussion) received anesthesia and incision equivalent to the corresponding preconditioned impact timing. Following impact or sham procedure, we sutured the mouse’s scalp closed and placed the mouse in a heated recovery cage until they exhibited normal signs of alertness and ambulation. Following recovery, animals were returned to the colony.

#### Brief vs Extended Impact Window Preconditioning

To determine the effect of the impact window on the cognitive outcome of subconcussive preconditioning, we evaluated two cumulative impact windows: 2 minutes (brief) and 48 hours (extended). Prior works show that impact timing is a key factor influencing the severity of repetitive impacts in both humans and rodents.^6,7,26^ We selected 3 total impacts in each window to minimize the detrimental effect of repetitive impacts (Fig 1 A), and to be consistent with prior work examining protective subthreshold preconditioning of concussion.^27^ We impacted the mice in the same location each time to ensure a spatially localized effect (Fig 1 B). For brief preconditioning, we administered 2 subconcussive impacts followed by a concussive impact, each of which were spaced a minute apart. For extended preconditioning, we administered the same impact profile but spaced the impacts 24 hours apart. For sham and unconditioned concussion for the extended window, we performed sham procedures 24- and 48-hours prior to the final sham or concussive procedure. Experimental Day 0 for both groups was the day of the final impact (Fig 1 C).^26^ We performed open field test 3 days post-injury (dpi), novel object recognition 3-4 dpi, and contextual fear conditioning 8-9 dpi. Following completion of behavioral testing 9 dpi, we sacrificed the animals for histology.

### Neurocognitive assessment

#### Open Field Test

At 3 days post injury (dpi), we placed individual mice in an open field chamber (12”x15”) containing no objects and allowed them to naturally explore for 15 minutes. We analyzed how much time the animal spent in corner, outer, inner, and center areas, as well as total ambulation using previously described automated methods.^52^ Open field test allows us to evaluate for motor deficits and anxiety behavior as anxious mice engage in thigmotaxis and remain at the perimeter of the field to avoid open spaces.

#### Novel Object Recognition

On Day 3 post-injury, we habituated the animals to the open field chamber for 15 minutes as described in *Methods: Open Field Test.* We then lightly cleaned the open field chamber to remove odor cues. Next, during training, we placed two identical objects in the open field and placed the same mouse in the chamber for two 10-minute trials to familiarize themselves with the objects, lightly cleaning between trials as before. On Day 4 post-injury, we replaced one of the familiar objects with a novel object in the same position. We placed the animal in the chamber to explore each object for a total of 5 minutes. We evaluated the time spent interacting with each object using automated methods described in Patel et al., 2014.^52^ Because mice inherently prefer novel objects, increased interaction with the novel object indicates recollection of object familiarity. Novel object preference refers to the time spent interacting with the novel object divided by the total time spent interacting with either object. Novel object recognition was performed by the same experimenter during the light cycle under similar temperature and lighting conditions. We excluded mice with an interaction time with either object less than 10 seconds total.^53^

#### Contextual Fear Conditioning

At 8 dpi, we placed the mouse in a foot shock chamber with sound attenuation for 3 minutes before delivering three shocks (0.4 mA) through the grid floor for a 2-second duration, 1 minute apart. One day later, at 9 dpi, we returned mice to the test chamber and recorded the animal movement over a 3-minute span. We removed odor cues with 10% bleach and 70% ethanol between trials. An experimenter blinded to experimental conditions manually scored freezing behavior from the video footage, defining fraction freezing time as the total accumulated freeze time over the total recording time. Contextual fear conditioning was performed by the same experimenter for all animals during the light cycle under similar temperature and lighting conditions. We excluded mice which were noted by the experimenter to exhibit fear and/or anxiety behaviors (including escape behaviors, increased defecation, etc) prior to training.^54^

### Histology

#### Perfusion and cryosectioning

We perfused mice at 1 and 9 dpi for immunohistochemistry as follows. We anesthetized the mice (250 mg/kg sodium pentobarbital, I.P.) and transcardially perfused them with ice-cold 0.9% saline (100 mL), followed by 300 mL of 4% paraformaldehyde (PFA). After 24 hours, we replaced the 4% PFA with 15-30% sucrose, and once the brain sank (another 24-48 hours later), we replaced with 30% sucrose again. We froze the brains in optimal cutting temperature (OCT) compound and sectioned the brain in 20 μm coronal sections approximately -1.5 to -2 mm from bregma.

#### Staining and fluorescence imaging

We stained 3 nonconsecutive sections/brain for 3 brains/condition. We blocked sections with Fish Serum Blocking Buffer (Thermo Scientific). For primary antibodies, we incubated sections with rabbit anti-Iba1 (1:200; Wako Chemicals) for 1 hour at room temperature and washed with 1X TBS with 0.1% Tween-20. Iba1 is expressed by both microglia and infiltrating macrophages. For secondary antibodies, we incubated with AlexaFluor 488 goat anti-rabbit (1:200; Abcam) for 30 minutes at room temperature and washed with 1X TBS with 0.1% Tween-20. We then incubated with NeuroTrace 640/660 Nissl Stain (1:30, Invitrogen), following manufacturer’s instructions to stain neurons. We mounted using ProLong Gold (Invitrogen). We imaged using a BZ-X800 Microscope (Keyence) at 20X magnification. We acquired images in a 10 μm z-stack with a 2 μm pitch, and then stitched together the best focused images using the BZ-X800 Analyzer Software (Keyence).

#### Image processing and stereology

For each image, we used ImageJ to examine the ipsilateral cortex directly below the impact site (approximately 1.5 mm^2^; Fig 3 G) and the ipsilateral hippocampus.

**Fig 3.**
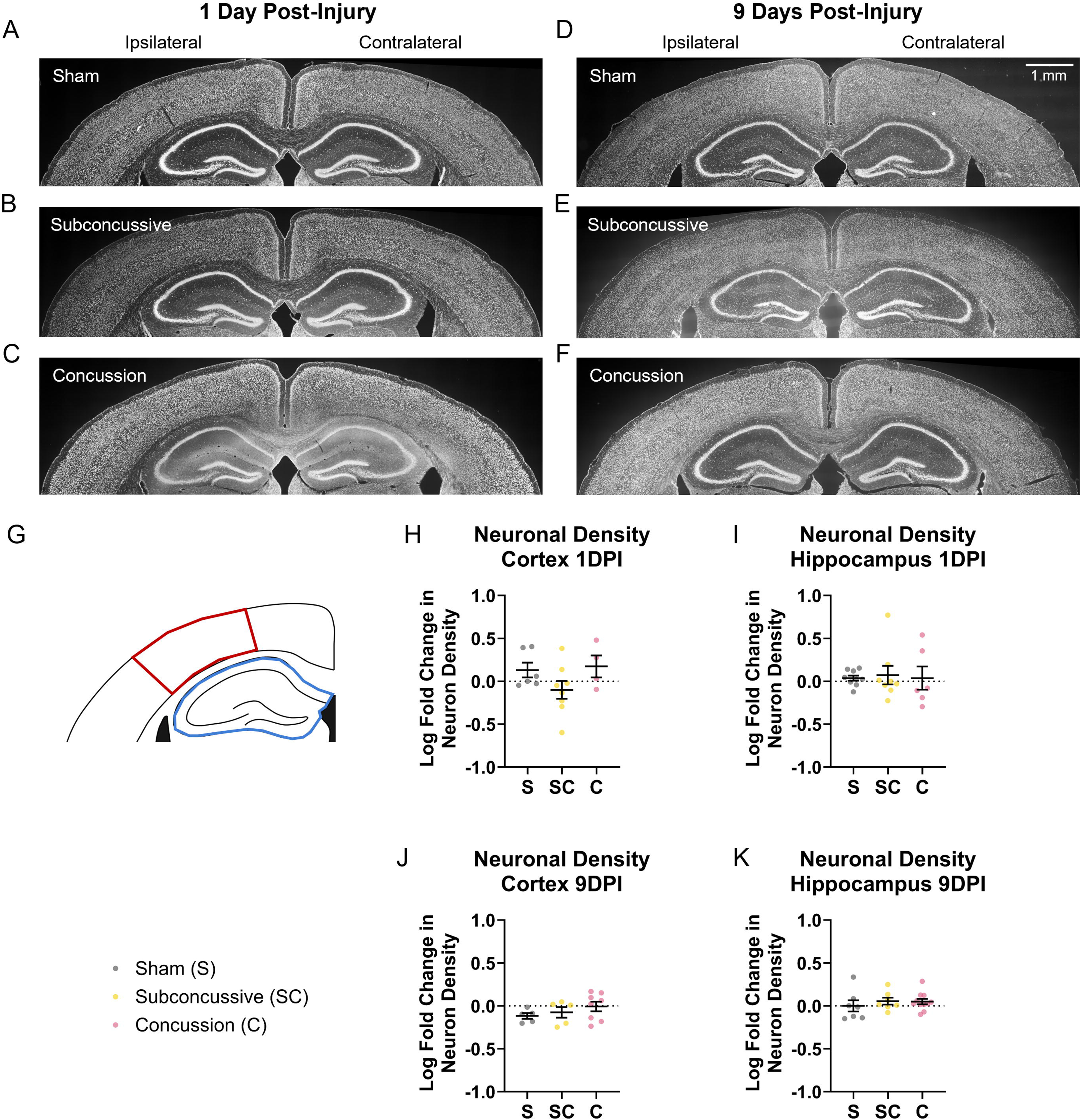
Neither subconcussive nor concussive impacts affect neuronal density. Representative image of NeuroTrace staining for (A-C) 1 day post injury (dpi) and (D-F) 9 dpi. (G) Areas of interest – cortex (red) and hippocampus (blue). (H-I) No change in ipsilateral versus contralateral neuronal density at 1 dpi in (H) cortex or (I) hippocampus, or (J, K) 9 dpi. Statistical testing was performed using linear mixed-effects models followed by Tukey’s post-hoc test. α = 0.05. Raw data shown as mean ± SEM. Left is ipsilateral; right is contralateral. S: Sham; SC: Subconcussive; C: Concussion.

To evaluate the severity of our head injuries, we estimated the changes in neuronal density in the cortex and hippocampus both 1 day and 9 days post injury. To estimate neuronal density, we enhanced the contrast on each image by equalizing the contrast histograms to account for variability in staining across sections, sharpened the image, identified regions of interest, applied a fixed threshold, and reported the percent area of the region of interest above threshold. We systematically excluded sections which had either high background or low staining efficiency in either hemisphere. We examined the fold change in the neuronal density in the ipsilateral hemisphere relative to the contralateral hemisphere to account for variability in staining across sections. Due to the density of neurons in the hippocampus, we elected to use this method rather than counting individual neurons.

To understand the morphological changes exhibited by microglia in response to injury, we counted and traced microglia in the ipsilateral cortex and hippocampus at both 1 and 9 dpi. To prepare images for counting and tracing, we applied a Fast Fourier Transform Bandpass Filter and adjusted the contrast on each image by equalizing the histograms as before. Again, we systematically excluded sections which had high background or blurriness which precluded confident counting and tracing. A scorer blinded to experimental condition manually counted all the microglia in a given region of interest. After manual counting, 30 microglia were randomly assigned for tracing from the total identified in a given region. For tracing, the scorer manually outlined the cell soma and traced its processes for Sholl analysis. Based on the traced soma, our software automatically identified the centroid and then drew concentric circles with radii increasing in 5 μm increments ^55^ for Sholl Analysis using the Sholl Analysis Package for ImageJ.^56^

### Statistical analysis

Microglia and neuronal statistical analysis were performed using Linear Mixed-Effects analysis in R Version 4.4.1 with Tukey’s post-hoc test to account for using multiple microglia from the same animal and for using multiple sections per animal.^57^ Behavioral statistical analysis was performed using one-way ANOVA (Novel Object Recognition, Contextual Fear Conditioning) or two-way ANOVA (Open Field Test) using Tukey’s method for multiple comparisons in GraphPad Prism Version 10 for Windows. Results are expressed as mean ± standard error of the mean (SEM) of the raw data.

## Results

### Head injury model captures severity-dependent cognitive deficits

First, we evaluated the neurobehavioral and neuronal response to our injury model. At our concussion impact level (3.5 m/s, 1 mm depth), there was no change in exploratory behavior or ambulation in the open field at 3 days post-injury (dpi; Fig 2 A; Suppl. Fig. 1 A), a significant deficit in novel object recognition at 4 dpi (Fig 2 B), and no contextual fear conditioning deficit at 9 dpi (Fig 2 C). Using scaling principles described in the Methods, our subconcussive impact (1 m/s, 1 mm depth) led to no significant impairment in open field behavior (Fig 2 A), novel object recognition (Fig 2 B), or contextual fear conditioning (Fig 2 C). There was no loss in neuronal density at either 1 or 9 dpi for either concussive or subconcussive impact in either the ipsilateral cortex or hippocampus relative to the contralateral side (Fig 3).

### Injury severity modifies timing of microglial hyperramification post-injury

Next, we evaluated the microglial response to our concussive and subconcussive impacts. To characterize this response, we examined the microglial number and morphology at 1 dpi to capture their early changes (Fig 4), and 9 dpi to measure more sustained changes in microglia response (Fig 5). Following concussion, microglia exhibited increased ramification (as measured by Sholl Analysis) at 1dpi in the hippocampus, indicating they assumed features of a hypersurveillant morphology (Fig 4 J). In contrast, at 9 dpi, the microglial morphology after concussive impacts was no different from sham (Fig 5 I). There were no other changes in response to concussion (Fig 4 E-I; Fig 5 D-H). Similarly, for subconcussive impacts, there was no change in the number of microglia or their soma size in the ipsilateral cortex at either 1 dpi or 9 dpi (Fig 4 E-I, Fig 5 D-H). However, at 9 dpi, but not 1 dpi, microglia from subconcussed brains exhibited increased ramification in the hippocampus (Fig 4 J; Fig 5 I). Taken together, these findings suggest that although concussed mice experienced a transient early increase in microglial ramification in the hippocampus, subconcussive impacts exhibited a more delayed increase in ramification.

**Fig 4.**
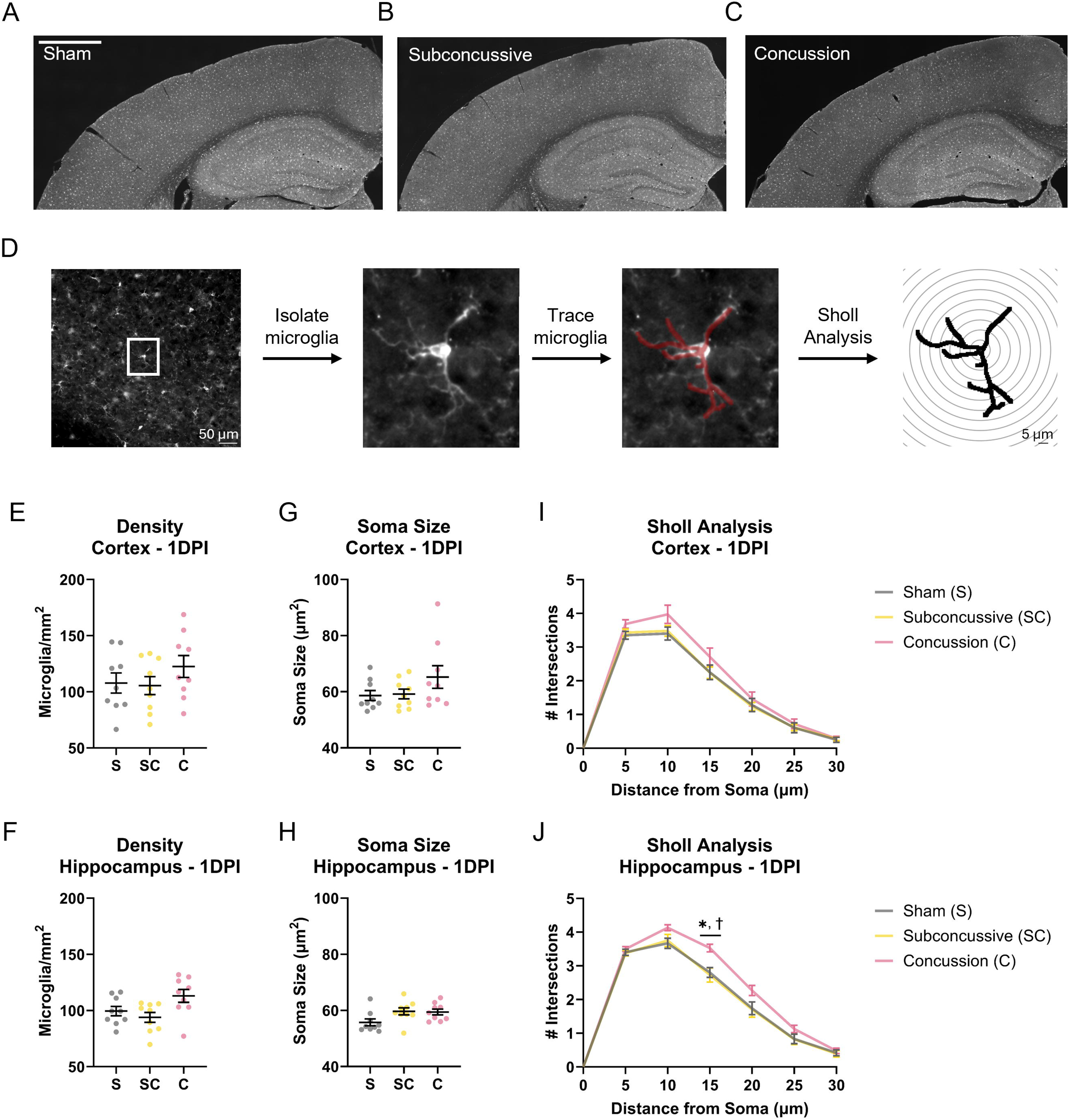
Microglia exhibit acute changes in morphology in the hippocampus in response to concussion. (A-C) Representative images of Iba1+ staining in ipsilateral cortex and hippocampus 1 day post-injury (dpi) (A) sham, (B) subconcussive impact, and (C) concussion. Scale bar = 1 mm. (D) Workflow for tracing microglia for Sholl analysis. (E, F) Microglia density in (E) cortex and (F) hippocampus, (G, H) soma size, and (I, J) Sholl Analysis. * indicates p<0.05 for concussion versus sham, † indicates p<0.05 for concussion versus subconcussive impact. Comparisons were performed using linear mixed-effects models with Tukey’s post-hoc test. Raw data shown as mean ± SEM. S: Sham; SC: Subconcussive; C: Concussion.

**Fig 5.**
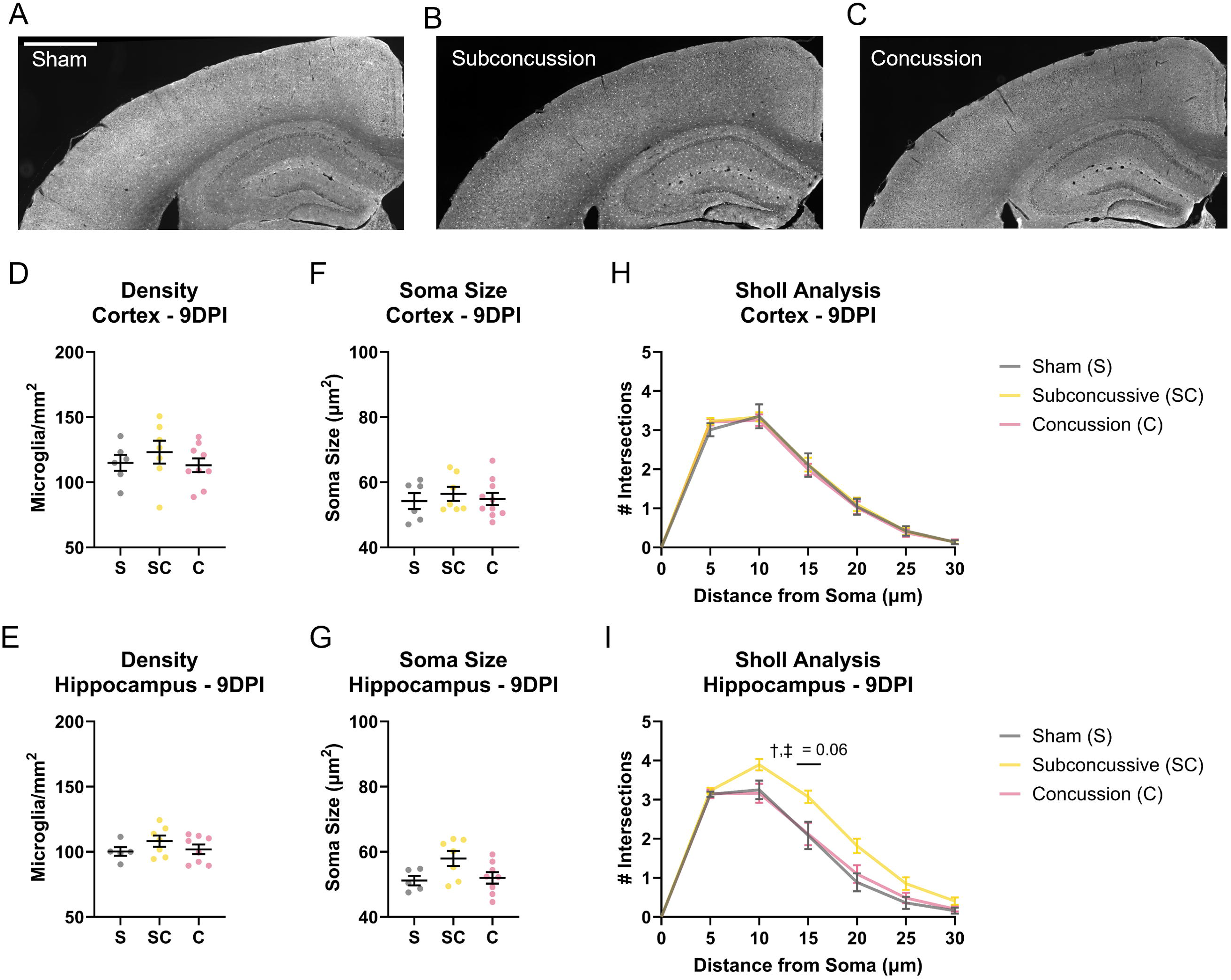
Microglia exhibit delayed changes in morphology in the hippocampus in response to subconcussive impact. (A-C) Representative images of Iba1+ staining in ipsilateral cortex and hippocampus 9 days post-injury (dpi) for (A) sham, (B) subconcussive impact, and (C) concussion. (E, F) Microglia density in (E) cortex and (F) hippocampus, (G, H) soma size, and (H, I) Sholl Analysis. ‡ indicates subconcussive impact versus sham, † indicates subconcussive impact versus concussion. Comparisons were performed using linear mixed-effects models with Tukey’s post-hoc test. Raw data shown as mean ± SEM. S: Sham; SC: Subconcussive; C: Concussion. Scale bar = 1 mm.

### Brief subconcussive preconditioning has no effect on microglial number or morphology

The more prolonged changes in microglial morphology which occurred after a subconcussive impact could be the basis of immune training or tolerance in a repetitive injury model, consistent with past reports in other disease models.^32,58^ We explored this possibility more thoroughly by examining whether the timing of subconcussive impacts could further shape these microglial responses. We hypothesized that two subconcussive impacts delivered prior to a single concussive impact over a brief 2-minute impact window would result in immune tolerance. Unlike either subconcussive or concussive impacts, there was no change in number of microglia, their soma size or ramification at 1 dpi in either the cortex or hippocampus relative to sham (Fig 6 A-F) or at 9 dpi (Fig 6 G-L). There was, however, a significant decrease in ramification in the hippocampus at 1 dpi relative to concussion (Fig 6 F). As before, there was no change in neuronal density in the cortex or hippocampus, at 1 dpi or 9 dpi, despite the increased impact load (Suppl. Fig. 2).

**Fig 6.**
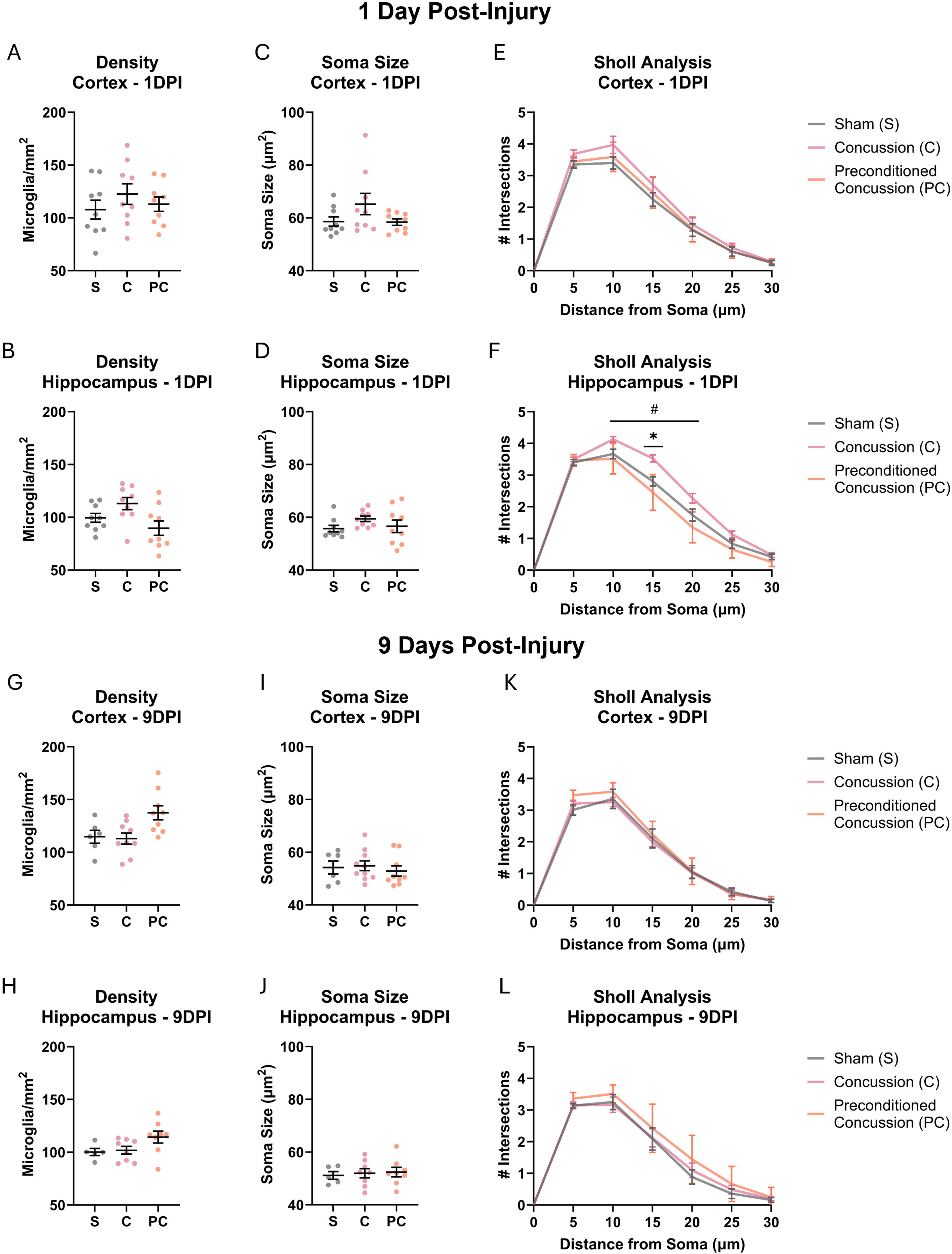
Brief subconcussive preconditioning prevents acute and delayed changes to microglia density and morphology. At 1 day post-injury (dpi), subconcussive preconditioning exhibits no change in (A, B) microglia density, (C, D) soma size, or (E, F) Sholl analysis. Similarly, at 9 dpi, subconcussive preconditioning also exhibits no changes in microglia (G, H) density, (I, J) soma size, or (K, H) Sholl analysis. Sham and concussion data are the same as reported in Fig. 4 and 5. * indicates p<0.05 for concussion versus sham; # indicates p<0.05 for concussion versus preconditioned concussion. * indicates p<0.05. Statistical testing was performed using linear mixed-effect models followed by Tukey’s post-hoc test. Raw data are presented as mean ± SEM. S: Sham; C: Concussion; PC: Preconditioned Concussion.

### Brief preconditioning, but not extended preconditioning, prevents concussion-associated cognitive deficits

We next considered whether the timing of subconcussive exposures would affect the resulting neurological impairments which normally occur in our closed head impact model. In human studies, there is a clear record of several subconcussive impacts that precede an impact causing concussion,^2–7^ but the relative effect of these cumulative exposures on cognitive outcomes is unknown. We found that both brief and extended preconditioning, like concussion, had no effect on exploration or ambulation in an open field at 3 dpi (Fig 7 A; Suppl. Fig. 1 B). However, brief preconditioning reversed concussion-associated deficits in novel object recognition at 3-4 dpi (Fig 7 B) and nearly significantly improved fear response relative to concussion at 8-9 dpi (Fig 7 C). These results suggest that brief preconditioning was able to improve cognitive outcomes across several domains (memory, anxiety) in the acute (3 days) and subacute (9 days) post-injury phase (Fig 1 C). Moreover, extended preconditioning did not create new deficits: neither exploration (Fig 7 D) nor ambulation were significantly changed (Suppl. Fig. 1 C). The rescue of cognitive deficits from preconditioning was related to the duration of the subconcussive response - extending the same two preconditioning impacts over 2 days, rather than 2 minutes, did not rescue concussion’s novel object recognition deficit (Fig 7 E) or fear conditioning (Fig 7 F).

**Fig 7.**
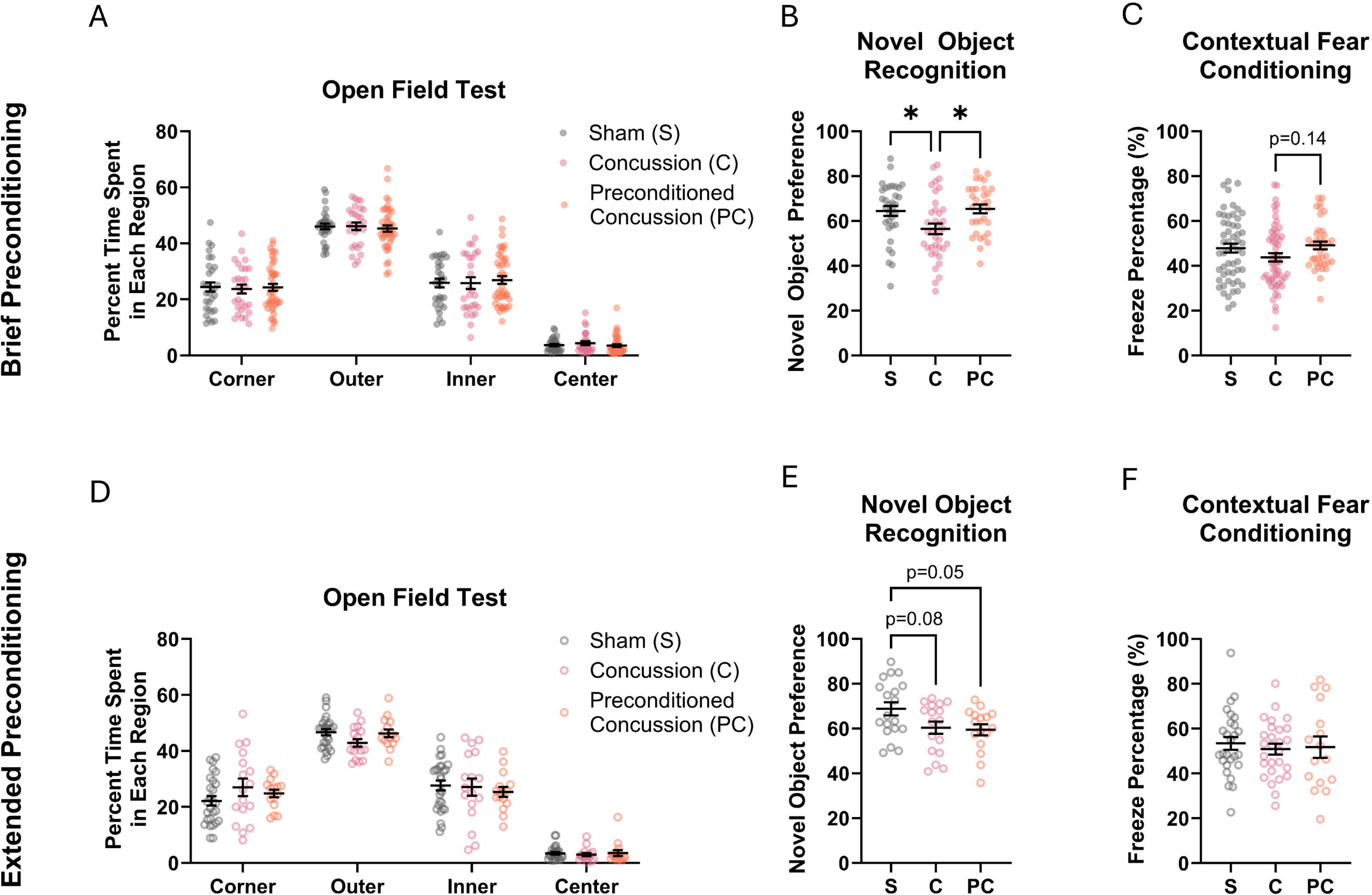
Brief, but not extended, preconditioning improves cognitive outcomes. (A) Brief preconditioning (2-minute window) had no effect on percent time spent in the corner, outer, inner or center areas of the open field. (B) Brief preconditioning significantly improves novel object preference compared to concussion. (C) Brief preconditioning improves fear memory relative to concussion, albeit non-significantly (p=0.14). Sham and concussion data in A-C are a reproduction of Fig 2. (D) Extended preconditioning (48-hour time window) had no effect on percent time exploring the open field. (E) Extended preconditioning produces the same near-significant deficit in novel object preference as concussion. (F) Neither concussion nor extended preconditioning significantly affects fear memory. * indicates p<0.05. Statistical testing was performed using two-way ANOVA (A, B) or one-way ANOVA (C-F) with Tukey’s post-hoc test. Data are shown as mean ± SEM. S: Sham; SC: Subconcussive; C: Concussion; PC: Preconditioned Concussion.

## Discussion

In this study, we examined how microglia increase ramification in response to different closed head impacts and, in parallel, how different sequences of head impacts can affect cognition. Consistent with past work, microglia display morphological changes early following concussion. These microglia changes occurred in parallel with impairments in novel object recognition and fear conditioning. In contrast, a single subconcussive impact produced a more delayed increase in ramification with no changes in neurobehavior. Rapidly delivering multiple subconcussive impacts before a concussive impact - one form of preconditioning - not only reduced changes in microglial morphology, but also erased the cognitive deficits associated with concussion. Using subconcussive impacts to reverse the deficits of concussion were time sensitive: extending these subconcussive impacts over days eliminated any protection offered by subconcussive preconditioning.

Previous studies have observed transient changes in microglia number and morphology following concussive and subconcussive impacts. In the first 72 hours following concussion, microglia increase in number at the injury site ^24,59,60^ and in soma size.^59,61^ These changes largely resolve within a week or two as the microglia decrease in number and return to a ramified surveillant state,^24,61^ but some other groups have observed more persistent increases in microgliosis.^60^ Our model produced changes on a similar timeline: the most profound changes in microglial morphology occurred one day after injury and resolved 9 days after the concussive injury. However, rather than an amoeboid morphology, we observed an increase in ramification typical of hypersurveillant or primed morphology.^62^ Other groups have noted this increase in ramification in response to concussion; for example, White and Vandevord observed a decrease in microglia and a simultaneous increase in branching 7 days post-injury (dpi).^63^ The heterogeneity in microglial response may be attributable to the variety in injury conditions and their severity. The effect of subconcussive impacts on microglia is less precisely characterized in past work. For both single and repetitive subconcussive impacts, some groups report a similar trend to concussion with an acute increase in the number of microglia that resolves within a week.^24,28^ Stemper and colleagues observed an increase in number and branching of microglia in response to highly repetitive subconcussive impacts at 14 dpi, suggesting that sufficient loading can produce persistent microglial pathology, potentially as severe as the acute concussive response we observe.^19^ Others have observed rod-like microglia in response to diffuse TBI, especially at acute and sub-acute timepoints,^64^ but we did not consistently observe this rare morphology in our regions of interest. We also did not observe amoeboid microglia at either time point, but these changes are frequently associated with phagocytosis in areas of necrotic cell death or tissue loss, neither of which we observe in our model.^65^ The more subtle changes we observe may be attributable to our less severe subconcussive impact.

Microglia can play a key role in learning and memory in both healthy and diseased states.^66^ For this reason, microglia are candidate mediators for the reversal of cognitive deficits we observed after subconcussive preconditioning. In healthy brains, for example, microglial production of TNF-L is required for sustaining fear memory,^67^ and microglial phagocytosis and synapse elimination is required for eliminating fear memory.^68^ Indeed, microglial activation during sensitive periods of development can impair neuron-microglia communication and synaptic modulation to produce learning deficits that only emerge days after the activation.^69^ In TBI, depleting microglia prior to and immediately following TBI is able to reverse cognitive deficits associated with TBI, suggesting that preventing acute inflammation may be an effective strategy for improving outcomes.^70,71^ Furthermore, cognitive deficits are associated with reactive microglial morphologies in TBI ^70,72^ and beyond.^73–75^ Taken together, our microglial analysis and neurocognitive analysis suggests that early increases in microglia with hypersurveillant morphology are associated with worse cognitive outcomes.

To our knowledge, there is no precedent for examining the microglial response to subconcussive preconditioning in a model of traumatic brain injury (TBI). Indeed, studies which study repetitive subconcussive impacts have largely focused on neuronal damage as a mechanism, e.g., as synaptic dysfunction ^18,21^ or white matter degeneration.^16^ Furthermore, those studies did not examine the interplay between subconcussive impacts and concussion outcomes. However, there are examples in similar neurological diseases, including ischemic stroke and epilepsy, which examine subthreshold preconditioning and microglial tolerance. In ischemic stroke, brief ischemia can be used to precondition against more prolonged ischemic events, and is associated with distinct microglia morphological changes and extent, much like what we observe with subconcussive impacts and preconditioning. Alternatively, kindling, a form of repetitive low-level seizure simulation, can be used to both lower the seizure threshold and induce spontaneous seizures^76^ and, less commonly, precondition the brain against seizures.^42,43,77^ Indeed, microglia have a beneficial role in secondary seizures and reduce the burden of kindled seizures ^78^ and kindling preconditioning regulates microglial gene expression.^77^ There is also a significant amount of overlap between these neurological diseases - e.g., ischemic preconditioning is effective in TBI ^79–81^ and TBI can induce posttraumatic epilepsy ^82^ and make the brain more susceptible to kindling ^38^ via neuroimmune mechanisms. Beyond subthreshold neurological stimulation, others have used systemic methods such as lipopolysaccharide (LPS) administration, exercise, heat, and anesthesia to confer neuroprotection against acute brain injury, and have identified changes in brain’s inflammatory response as a key mechanism.^83^ Most common among these is LPS preconditioning, which directly and systemically stimulates the body’s innate immune response.^84^ At low doses, LPS is protective against subsequent TBI, reducing cell death and neurodegeneration, improving neurocognitive function, reducing pro-inflammatory cytokines, and reducing phagocytic microglia.^33–35^ However, a key challenge with using LPS for neuroprotection is its systemic effects, including acute weight loss^33^ and locomotor impairments.^34^ Together, this past work linking the response of microglia to the neurological outcome has led to a broad rationale for identifying brain-specific immunological therapeutic strategies.

In addition to desensitizing the brain prior to a more severe injury, microglia can play several roles in the brain during recovery. Microglia play a key role in synaptic remodeling,^85^ and hyperramified microglia in particular are hypothesized to mediate dendritic spine loss.^86^ One way microglia can modify synapses is by remodeling the extracellular matrix,^87^ including the perineuronal nets which stabilize synapses ^88^ and may underpin long-term memory storage.^89^ Microglia also secrete factors such as cytokines and other neurotrophic factors which can affect inflammatory cascades and synaptic plasticity.^90^ In preliminary studies, we did not observe long-lasting changes in a panel of serum cytokine levels (data not shown), but that does not rule out earlier changes. Microglia also remove cellular debris following injury.^90^ However, given the absence of neuronal injury or tissue loss in our injury model, as well as the absence of amoeboid microglia which are often associated with phagocytosis,^62^ we do not expect this to be a significant microglial role affected by our model. It is more likely that preconditioning affects microglia-mediated synaptic plasticity to improve cognitive function, perhaps through the delayed appearance of the hyperramified microglia after subconcussive preconditioning.

In summary, our work shows that rapidly delivered subconcussive impacts can precondition the brain against the deficits normally caused by concussion. We find that changes in microglial morphology differ between subconcussive and concussive impacts, and differ yet again with preconditioning. In general, the alterations in microglial morphology resemble similar changes reported in other models of preconditioning. As evident through past work in preconditioning, our initial observations provide the groundwork for exploring more directly the role of microglia and, in the longer term, revealing the mechanism(s) which affect the timing and duration of this phenomenon. Together, these findings can provide critical insights into expanding the role of microglia from surveillance and maintenance in the injured brain into providing temporary protection of the brain to damaging insults.

### Transparency, Rigor and Reproducibility Statement

All surgical procedures were performed in accordance with the IACUC at the University of Pennsylvania. A sample size of 3 mice/experimental condition with 3 sections/brain was planned based on prior work.^91^ For microglia analysis, an experimenter blinded to the experimental condition counted all microglia in a region of interest and then randomly assigned 30 microglia for morphological analysis, similar to prior studies.^55^ A mixed-effects model was selected for analysis of microglial soma size and Sholl analysis to account for variability in sections from the same brain.^55,57^ A sample size of 25 mice/experimental condition was planned for behavioral studies to detect significant differences with L=0.05 and 80% power. However, as this is a new TBI model, the sample size was increased to account for greater-than-anticipated variability in outcomes. Exclusion criteria can be found in the Methods. Novel object recognition and contextual fear conditioning were performed during the light cycle by the same experimenter under similar conditions (e.g., time of day, temperature, lighting). Behavioral analysis was conducted by an experimenter blinded to the experimental condition. Multiple comparisons testing was performed using Tukey’s method.

## Acknowledgements

We thank D. Kacy Cullen for the use of Keyence BZ-X800 for fluorescence imaging, and thank Jared A. Rifkin and Melanie C. Hilman for editorial feedback.

## Authorship Contribution Statement

E.D.A.: conceptualization, methodology, software, formal analysis, investigation, writing - original draft preparation, writing - review & editing, visualization, validation, data curation

K.K.: formal analysis, investigation, software, validation, writing - review & editing

A.P.G.: methodology, formal analysis, validation, project administration

A.N.: methodology, formal analysis, software

E.G.: methodology, formal analysis, software, validation, data curation

D.V.A.: methodology

D.F.M.: supervision, conceptualization, methodology, funding acquisition, writing - review & editing

---

**Suppl. Fig. 1. Ambulation in an open field is unaffected by our injury model.**

Neither (A) subconcussive, (A, B) concussive, (B) brief preconditioning, or (C) extended preconditioning affects ambulation in an open field 3 days post injury. Statistical testing performed using one-way ANOVA with Tukey’s post-hoc test. Data presented as mean ± SEM. α = 0.05. S: Sham; SC: Subconcussive; C: Concussion; PC: Preconditioned Concussion.

**Suppl. Fig. 2. Subconcussive preconditioning has no effect on neuron density 1 or 9 days post-injury.**

Representative image of NeuroTrace staining for subconcussive preconditioning (A) 1 day post injury (dpi) and (B) 9 dpi. No change in ipsilateral versus contralateral neuronal density 1dpi in (C) cortex or (D) hippocampus, or (E,F) 9 dpi. Statistical testing was performed using linear mixed-effects models followed by Tukey’s post-hoc test. α = 0.05. Left is ipsilateral; right is contralateral. S: Sham; C: Concussion; PC: Preconditioned Concussion.

