## Supplementary figures and images for "Subconcussive preconditioning prevents microglial morphology changes and improves cognitive outcomes in mice"

### Supplemental Figure 1

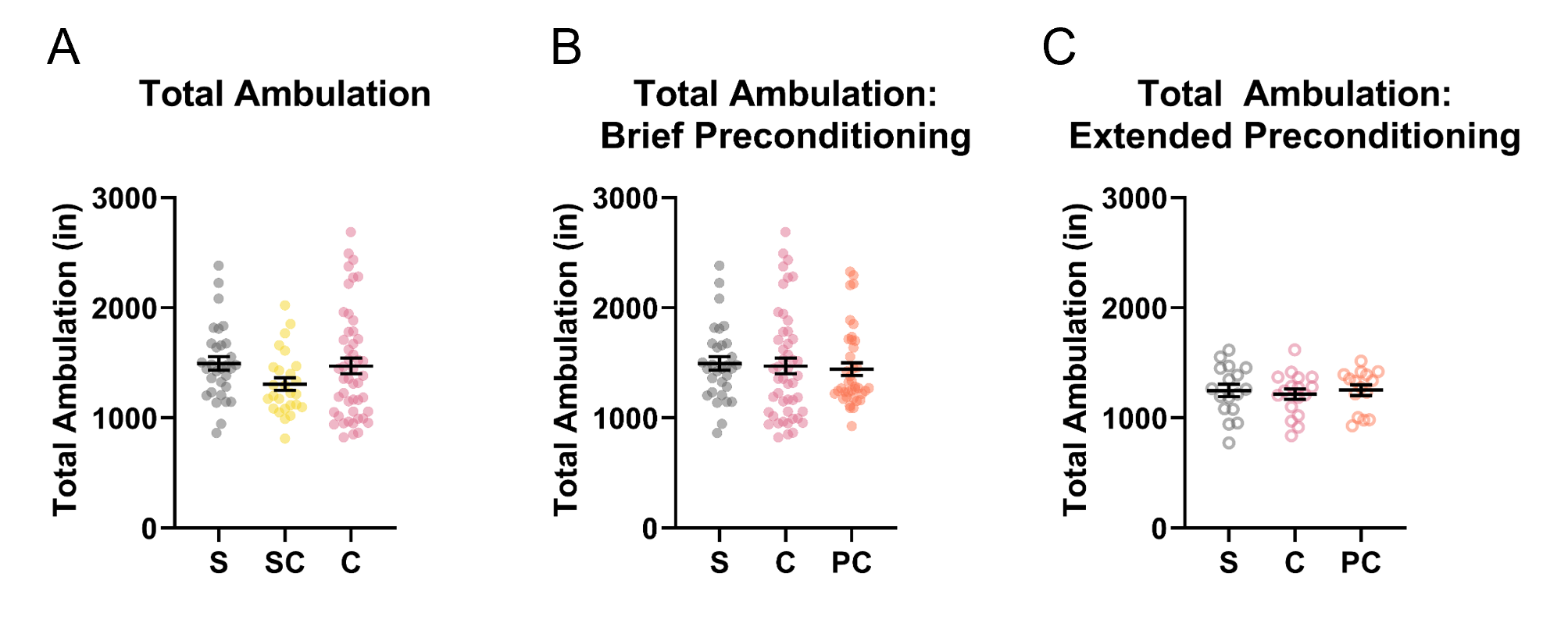

### Supplemental Figure 2

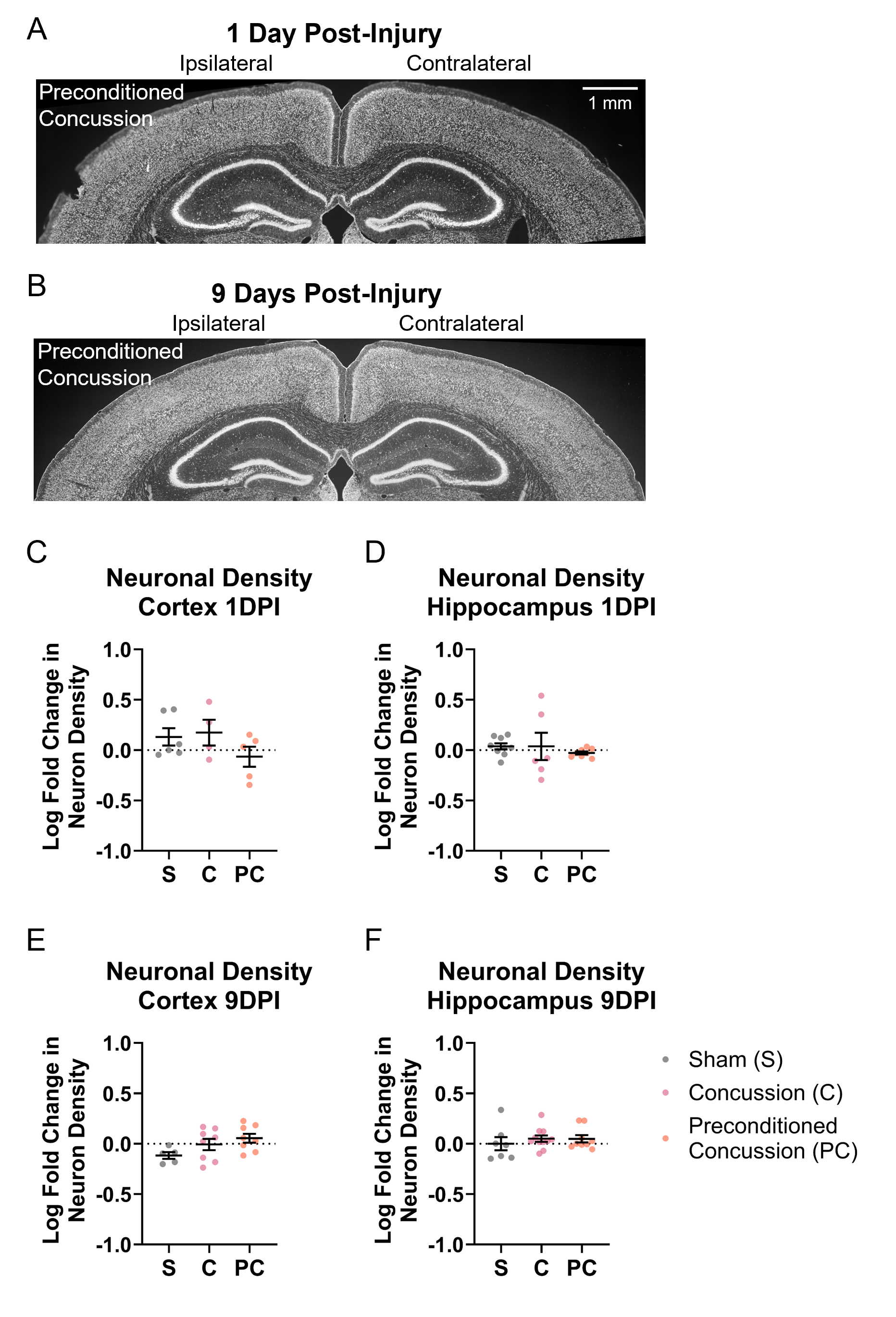
